# Multisensory processes can compensate for attention deficits in schizophrenia

**DOI:** 10.1101/2020.08.14.251405

**Authors:** James K. Moran, Julian Keil, Alexander Masurovsky, Stefan Gutwinski, Christiane Montag, Daniel Senkowski

**Author notes:** **Data availability statement**: Data not available due to ethical restrictions. **Ethics approval statement:** The study was carried out in accordance with the 2008 Declaration of Helsinki and was approved by the ethics commission of the Charité – Universitätsmedizin Berlin (Approval number:EA1/169/11). **Author Contributions:** Study Design: JK, DS, Data Collection: JKM, AM, SG, CM, Data Analysis: JKM, AM, DS; Manuscript preparation: JKM, JK, AM, SG, CM, DS.

## Abstract

Studies on schizophrenia (SCZ) and aberrant multisensory integration (MSI) show conflicting results. These divergent results are potentially confounded by attention deficits in SCZ. To test this, we examined the interplay between MSI and intersensory attention (IA) in healthy controls (N=27) and in SCZ (N=27). Evoked brain potentials to unisensory-visual (V), unisensory-tactile (T) or bisensory VT stimuli were measured with high density electroencephalography, whilst participants attended block-wise to either visual or tactile inputs. Behaviourally, IA effects in SCZ are uncompromised for bisensory stimuli, but diminished for unisensory stimuli. At the neural level, we observed reduced IA effects for bisensory stimuli over mediofrontal scalp regions (230-320ms) in SCZ. The analysis of MSI revealed multiple phases of integration over occipital and frontal scalp regions (240-364ms), with comparable performance between HC and SCZ. The magnitudes of IA and MSI effects were both positively related to the behavioural performance in SCZ, indicating that IA and MSI mutually facilitate bisensory stimulus processing. Our study suggests that widely intact MSI, which facilitates stimulus processing, can compensate for top-down attention deficits in SCZ. Further, the interplay of IA and MSI implies that differences in attentional demands may account for previous conflicting findings on MSI in schizophrenia.

---

Veridical understanding of the physical world requires coordination of information from different sensory modalities. This coordination involves selectively attending to some sensory stimuli and filtering others. To comprehend a multisensory object, sensory input has to be integrated across the different senses and this integration has been shown to capture attention (Talsma et al., 2010). There is some evidence for aberrant multisensory integration (MSI) in SCZ (Stevenson et al., 2017; Williams et al., 2010), but findings are controversial (e.g. Stone et al., 2011; Wynn et al., 2014). It may be that differences in top-down attention demands contribute to the inconsistent findings. Since there is robust evidence of unisensory attention deficits in SCZ (Berkovitch et al., 2018; Sauer et al., 2017), examining the interplay between intersensory attention (IA) and MSI may shed light on the inconsistency in findings regarding multisensory processing in SCZ.

There is a well-founded dependency between MSI and attention (Talsma et al., 2010). This is visible in early components of event-related potentials (ERPs) measured during IA tasks (Spence & Driver, 1997). In an EEG study, Lange and Röder (2006) found that orienting attention towards visual or tactile sensory stimuli in a multisensory task increases early negative deflections in primary visual and somatosensory cortices, respectively. Similarly, ERPs in a visual-tactile (VT) attention task are greater when stimuli are attended compared to when they are unattended (Keil et al., 2017). Thus, IA has measurable effects on multisensory processing, as reflected in evoked electrical brain activity.

Thus far, the available studies examining MSI in SCZ have revealed mixed results. For instance, Stevenson et al., (2017) found deficits in both unisensory and multisensory performance for temporal order judgment tasks. Similarly, in a basic audiovisual target detection task, some studies found deficits in both unisensory and multisensory performance in SCZ, relative to healthy controls (HC) (Williams et al., 2010). However, others found unisensory processing deficits in target detection in SCZ, but unimpaired MSI in both performance and ERP parameters (Stone et al., 2011; Wynn et al., 2014). Hence, there are consistent findings of unisensory processing deficits in SCZ, but much less consistent findings of deficits in multisensory tasks.

The centrality of attention to MSI is relevant to SCZ, as people with SCZ show deficits in variousfacetsofattention (Gold et al.,2018). Studies have found aberrant top-down processing in SCZ, as reflected in the P300 component (Bramon, 2004). Other studies have shown impairments in top-down visual search (Fuller et al., 2006; Gold et al., 2007) and performance in mismatch negativity tasks in situations where predicted sounds are omitted as well as with unpredictable sounds, suggesting weakened top-down predictive coding in SCZ (Sauer et al., 2017). Taken together, top-down attention deficits and their neurobiological correlates are established in SCZ, while findings of MSI deficits in SCZ remain controversial. In this study we adapted a visual-tactile attention paradigm (Keil et al., 2017; Pomper et al., 2015) to examine the interplay between MSI and IA in healthy control (HC) participants and in people with SCZ. Using a rigorous data-driven analysis approach, we compared the effects and interplay of IA and MSI on behavioural data and ERPs between HC and SCZ.

## Materials and Methods

### Sample and clinical data

Twenty-nine people diagnosed with SCZ according to the DSM-5 criteria, were recruited at the outpatient units of the Charité–Universitätsmedizin, Berlin. After exclusion of two behavioural outliers, 27 patients were included in the final analysis (see Table 1). Psychiatric assessment was undertaken by an experienced psychiatrist at the recruiting institution. Patients taking the following medications were excluded from the study, in order to minimize distorting effects upon the EEG (Aiyer et al., 2016):benzodiazepines, lithium, valproic acid, and haloperidol. Of 29 control participants recruited from the general population, two were excluded during the analysis of behavioural (N = 1) or EEG (N = 1) data. Hence, 27 control participants were included in the final data analyses. Study groups were matched for handedness(Oldfield, 1971), education, smoking, age, and gender (Table 1). Furthermore, they were screened for comorbid psychopathology using the German version of the Structured Clinical Interview for DSM-4-TR Non-Patient Edition (SCID). The cognitive capacity of all participants was assessed with the Brief Assessment of Cognition in Schizophrenia(BACS; Keefe et al., 2004). Symptom severity in SCZ was tested using the Positive and Negative Symptom Scale (PANSS; Kay, Fiszbein, & Opfer, 1987) and items were grouped according to the 5-factor model (Wallwork et al., 2012). All participants gave written informed consent, including awareness of EU data protection laws and had normal hearing and normal/corrected to normal vision. People with neurological disorders or previous head injury with loss of consciousness were not included in the study. Furthermore, in the control group, those with immediate family members with a psychiatric disorder or neurological disorder were excluded. All participants underwent a drug screening (Drug-Screen Multi 5 Test, Nal von Minden, Amphetamine, benzodiazepine, cocaine, opioids and cannabis) prior to measurement. The study was carried out in accordance with the 2008 Declaration of Helsinki and was approved by the ethics commission of the Charité– Universitätsmedizin Berlin (Approval number: EA1/169/11).

**Table 1:**
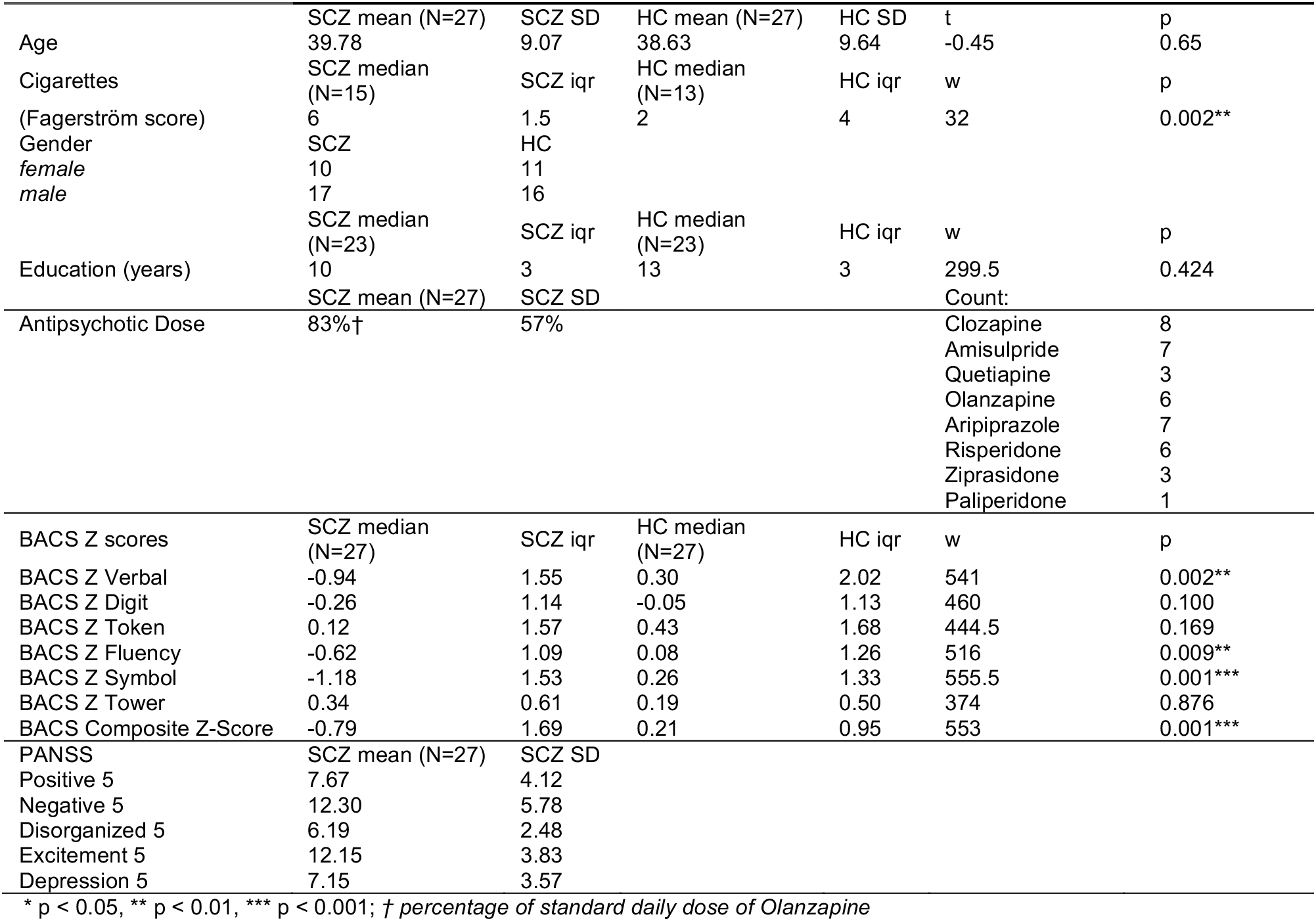
Demographics for participants. Differences between groups were calculated by either parametric independent t-tests (t) or non-parametric Wilcoxon tests (w), where data did not fulfil assumptions of parametric tests, iqr = inter-quartile ratio. Antipsychotic dose is calculated as the percentage of standard daily dose of Olanzapine, which is 10mg. BACS raw scores were converted to z-scores normalized for age and gender.; SD = standard deviation.

|  |  |  |  |  |  |  |
| --- | --- | --- | --- | --- | --- | --- |
| Age | SCZ mean (N=27)<br>39.78 | SCZ SD<br>9.07 | HC mean (N=27)<br>38.63 | HC SD<br>9.64 | t<br>-0.45 | p<br>0.65 |
| Cigarettes<br>(Fagerström score) | SCZ median<br>(N=15)<br>6 | SCZ iqr<br>1.5 | HC median<br>(N=13)<br>2 | HC iqr<br>4 | w<br>32 | p<br>0.002** |
| Gender<br>*female*<br>*male* | SCZ<br>10<br>17 | HC<br>11<br>16 |  |  |  |  |
| Education (years) | SCZ median<br>(N=23)<br>10 | SCZ iqr<br>3 | HC median<br>(N=23)<br>13 | HC iqr<br>3 | w<br>299.5 | p<br>0.424 |
| Antipsychotic Dose | SCZ mean (N=27)<br>83%† | SCZ SD<br>57% |  |  | Count: |  |
|  |  |  |  |  | Clozapine | 8 |
|  |  |  |  |  | Amisulpride | 7 |
|  |  |  |  |  | Quetiapine | 3 |
|  |  |  |  |  | Olanzapine | 6 |
|  |  |  |  |  | Aripiprazole | 7 |
|  |  |  |  |  | Risperidone | 6 |
|  |  |  |  |  | Ziprasidone | 3 |
|  |  |  |  |  | Paliperidone | 1 |
| BACS Z scores | SCZ median<br>(N=27) | SCZ iqr | HC median<br>(N=27) | HC iqr | w | p |
| BACS Z Verbal | -0.94 | 1.55 | 0.30 | 2.02 | 541 | 0.002** |
| BACS Z Digit | -0.26 | 1.14 | -0.05 | 1.13 | 460 | 0.100 |
| BACS Z Token | 0.12 | 1.57 | 0.43 | 1.68 | 444.5 | 0.169 |
| BACS Z Fluency | -0.62 | 1.09 | 0.08 | 1.26 | 516 | 0.009** |
| BACS Z Symbol | -1.18 | 1.53 | 0.26 | 1.33 | 555.5 | 0.001*** |
| BACS Z Tower | 0.34 | 0.61 | 0.19 | 0.50 | 374 | 0.876 |
| BACS Composite Z-Score | -0.79 | 1.69 | 0.21 | 0.95 | 553 | 0.001*** |
| PANSS | SCZ mean (N=27) | SCZ SD |  |  |  |  |
| Positive 5 | 7.67 | 4.12 |  |  |  |  |
| Negative 5 | 12.30 | 5.78 |  |  |  |  |
| Disorganized 5 | 6.19 | 2.48 |  |  |  |  |
| Excitement 5 | 12.15 | 3.83 |  |  |  |  |
| Depression 5 | 7.15 | 3.57 |  |  |  |  |

\* p < 0.05, \*\* p < 0.01, \*\*\* p < 0.001; † percentage of standard daily dose of Olanzapine

### Setup and Procedure

Participants were seated in an electrically and acoustically shielded chamber with low lighting. They were presented, in random order, with unisensory-visual (V), unisensory-tactile (T), and bisensory VT stimuli and had to detect occasional target stimuli in either the visual or tactile modality. Visual stimuli were presented for 150ms against a neutral gray background with a luminance of 30cd/m^2^ at the centre of a tilted TFT monitor (see Figure 1, left panel). The visual standard stimulus consisted of a Gabor patch in a circular frame, (diameter: 5.75°, spatial frequency = 1 cycle per degree, Gaussian standard deviation = 2°). The target visual stimulus was the same stimulus but flickering at 16.7Hz. The Braille stimulator was attached to the back of the monitor in the centre so that visual and tactile stimuli were spatially aligned. Tactile standard and target stimuli were administered by a piezoelectric Braille stimulator (QuaeroSys, St. Johann, Germany),consisting of 16 pins arranged in a square, with 2.5mm spacing between the pins. For the standard tactile stimulus, these were elevated onto the participant’s left index finger for 150ms. The target tactile stimulus consisted of multiple high frequency elevations and contractions at 16.7 Hz for 150ms. An auditory mask of white noise was presented during the experimental blocks to cancel out the sound of the Braille stimulator. The effectiveness of the white noise was verified prior to testing, by asking participants if they could hear anything during presentation of sample tactile stimuli.

The procedure began with presentations of samples of the different stimulus types. Participants performed a speeded response task by pressing a button with their right index finger when a target in the attended modality appeared. There were a total of 1722 trials presented across 14 blocks (i.e.,123 trials per block). Each block lasted about 4 minutes, alternating blockwise between visual and tactile attention tasks. There were 861 of both visual and tactile attention standard trials, broken down into 235V, 235T and two sets of 235 bisensory VT trials per attention condition. In addition, 52 unisensory V, 52 unisensory T and 52 bisensory VT target trials (where both sensory constituents were targets) were presented. Thus about 18% of trials overall were target stimuli. The stimuli were presented for 150ms, following this, participants were given 1000ms to respond (or to not respond in case of standards). The interstimulus interval (ISI) was randomized between 600 to 1000ms (average 800ms) between the stimulus presentation and response time. The response interval was indicated by a transformation of the fixation cross into a circle to cue the response (Figure 1, right panel).

**Figure 1:**
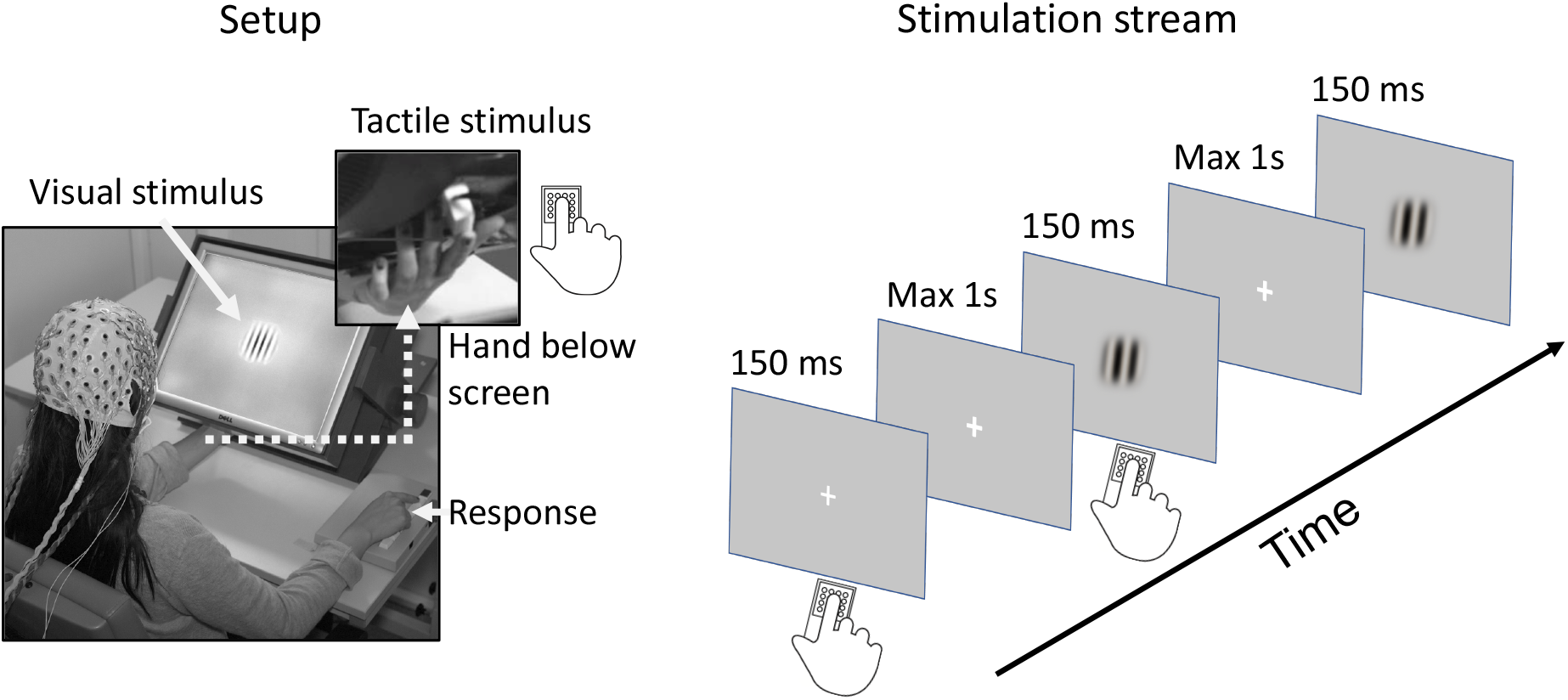
Outline of the visuo-tactile attention paradigm. Left panel: Illustration of the experimental setup. The monitor was tilted on an angle, and the participant’s hand was placed behind the screen to where the tactile stimulator was located. The left hand was held in place with cushioning to avoid muscle movement or fatigue. Tactile stimulation was presented to the left hand, and behavioural responses were given with the right-hand index finger. In the bisensory stimuli, the visual and tactile inputs were spatio-temporally aligned. Right panel: Illustration of the unisensory and bisensory stimulation stream.

Participants had to block-wise detect either visual (attend-visual condition) or tactile (attend-tactile condition) target stimuli that appeared occasionally. The inter-trial interval was randomized between 600 and 1000ms (mean 800ms).

### EEG Recording

Data were recorded using a 128-channel passive EEG system (EasyCap, Herrsching, Germany), which included two EOG electrodes (online: 1000 Hz sampling rate with a 0.016 – 250 Hz bandpass filter; offline: 49 – 51 Hz, 4th order Butterworth notch filter, 125 Hz 24th order FIR lowpass filter, down sampled to 500 Hz, 1 Hz 1500th order FIR highpass filter). Data was re-referenced to the average of all EEG electrodes. Non-stationary artifacts were identified by visual inspection, and the contaminated trials removed. After this process, there was no significant difference between remaining trials between groups (SCZ: M = 1481.67 [SD = 143.09] trials; HC = 1488 [SD = 173.58] trials; t(50.05) = 0.16, p = 0.870). Independent component (IC) analyses to correct for EOG and ECG artifacts were conducted, (Chaumon, Bishop, & Busch, 2015; Lee et al., 1999). The median number of components rejected was 3 for each group. Spherical interpolation was used to interpolate remaining noisy channels. There was no significant difference in the number of channels remaining (SCZ = 118.14 [SD = 5.25] channels; HC = 116.48 [SD=4.50] channels, t(51.18) = −1.27, p = 0.209). One HC was rejected at the preprocessing stage because of excessive eye-blinks, which were not removable by ICA.

### Analysis of behavioural data

To analyse hit-rates vs. false-alarm rates, d-prime values of behavioural responses were calculated (Stanislaw & Todorov, 1999). A successful hit was defined as a response to the correct target stimulus, e.g., in the visual attention condition, the participant pressed a response when a visual target appeared. The false response to this would be when a participant responds instead to a standard stimulus, e.g., for visual attention, the participant presses the button for a standard visual stimulus. Similarly, for the bisensory VT stimuli, a correct response (visual and tactile target) was contrasted with a false response to bisensory standard stimuli. Median percentage of responses outside of the analysis range of 100ms to 900ms were 0.05% for SCZ and 0.05 % for HC (Wilcoxon test, W = 264.5, p-value = 0.356). For the statistical analysis, a three-way ANOVA was conducted with the factors Group (SCZ vs. HC), Attended modality (tactile vs. visual), and Mode (bisensory vs. unisensory).

### Analysis of evoked brain activity

To avoid confounding motor activity and in line with previous studies on IA (Talsma et al., 2009; Talsma & Woldorff, 2005)only standard stimuli were included in the EEG data analysis. Moreover, EEG data were specifically examined for bisensory VT stimuli, because the current setup provided the unique opportunity to compare IA effects and MSI effects in the same bisensory stimuli. EEG data were analysed for the 50ms to 400ms poststimulus interval. The statistical analysis of IA and MSI effects was conducted separately and took place in two steps: (i) Data-driven clustering algorithms and permutation tests were used to define regions of interest (ROI) and times of interest (TOI) for the within-subject experimental manipulations (Maris & Oostenveld, 2007). As the clusters were defined both temporally and spatially, it was possible to have temporally overlapping clusters, e.g., a frontal and occipital ROI occurring more or less simultaneously, but with slight differences in temporal onset of significance. For the cluster analysis of IA, data were first combined across groups (SCZ and HC) and the difference between the two attention conditions was tested. For the cluster analysis of MSI, the additive approach was applied, whereby evoked responses to bisensory VT stimuli were compared with the linear combined brain responses to unisensory stimuli (i.e. V+T; Molholm et al., 2002). (ii) These within-subject contrasts of attention (Visual vs. Tactile) or MSI (bisensory vs. additive) were tested with dependent- samples *t*-tests with Monte–Carlo randomization and cluster-based correction for multiple comparisons (Maris & Oostenveld, 2007).(iii) The averaged amplitude of the clusters defined in the first step was then analysed in two-by-two factorial mixed model ANOVAs, which specifically focused on interactions in relation to the factor Group.

In order to enable interpretation of any null results, we computed Bayes Factors (BF10, Rouder et al.,2009)as an indicator of the relative evidence for the H0 and H1.BFs between 1–3 indicate anecdotal support for the alternative hypothesis (H1) while BF between 3–10 and above 10 indicate respectively moderate and strong support for H1. BF = 1 indicates equal support for H1 and null hypothesis (H0) while BF between 1/3–1, 1/10–1/3 and below 1/10, provide respectively anecdotal, moderate and strong support for H0 (Aczel et al.,2017).

The time windows in which main effects or interactions of the factor Group were found were correlated against behavioural performance in the SCZ group to further examine the relationships between IA, MSI, and behavioural performance in patients. Finally, correlations between PANSS and BACS and potential confounds (Olanzapine equivalent dose, Fägerstrom tests) and all significant clusters were calculated. All comparisons together were adjusted with Benjamini-Hochberg corrections (Benjamini & Hochberg, 1995).

## Results

### Behavioural data

The three-way ANOVA using the factors Group, Attended modality, and Mode revealed a significant main effect of Mode (F(1,156) = 49.27, p < 0.0001), showing that responses to bisensory VT stimuli were overall more accurate than those for unisensory stimuli (β = 0.18). There was also a main effect of Attended modality (F(1,156) = 13.02, p < 0.0001), demonstrating that attending to tactile stimuli produced more accurate responses than attending to visual stimuli(β = 0.22). No significant main effect of Group was observed (F(1,52)=3.74, p=0.059,β = −0.04). There was also a two-way interaction between Group and Mode(F(1,156) = 4.34, p = 0.039, β = 0.22). Follow-up simple contrasts between groups and averaged across attention conditions revealed significant group differences for unisensory stimuli(HC: adj-m= 4.43 [4.01-4.86], SCZ: adj-m=3.82 [CI: 3.39-4.25], t(69.5) = 2.57, p = 0.012, BF10 = 3.88)but not for bisensory stimuli (HC: adj-m = 4.87 [CI: 4.43-5.30], SCZ: adj-m=4.63 [CI: 4.20-5.06], t(69.5)=1.02, p=0.312,BF10=0.42).The contrast of the difference between these differences was significant(m=0.37,t(156)=2.08,p=0.039,BF10=1.59),indicating that HC performed better than SCZ for unisensory stimuli, but not for bisensory stimuli. Thus, on the one hand, our behavioural analyses demonstrate attention deficits for unisensory stimuli in SCZ, whereas there was evidence that performance on bisensory stimuli was comparable between HC and SCZ.

### Intersensory attention effects in evoked brain activity

The analysis of IA effects on bisensory VT stimuli across combined HC and SCZ groups revealed five significant clusters (Figure 3 and Table 2). The first cluster was right centrally localized (90-108ms). This is followed by two temporally overlapping clusters encompassing occipital (158-216ms) and frontal 168-218ms) regions, and two temporally overlapping later clusters in mediofrontral (230-320ms) and occipital (282-318ms) scalp regions. In the next step of the analysis, two-by-two factorial mixed model ANOVA were computed for each cluster. The main focus of these ANOVAs was on the group by attention interactions. For the first three clusters, no such interactions were found, indicating comparable effects of IA between group up to218ms. For the latter two clusters, encompassing mediofrontal and occipital brain regions, Group by Attention interactions were found (230-320ms β = −0.19, p = 0.025; 282-318ms β = 0.21, p = 0.002). The interactions were explained by the fact that the HC group showed larger ERPs in the attend visual vs. attend tactile condition for 230-320ms conditions (HC mean vis– tac=-0.44, t(52)= −5.77,p<0.0001, BF=27554.27;SCZ mean vis–tac= −0.19, t(52)= −2.52,p=0.015, BF10= 3.52). The difference between these two contrasts showed that HC had a larger difference between vis and tac conditions than SCZ (SCZ-HCm = −0.25 t(52)= −2.30 p = 0.026, BF10= 2.32).The same pattern of larger ERPs differences between attend visual vs. attend tactile was found in the 282-318ms cluster (HC mean vis – tac = 0.57, t(52) = 6.75, p < 0.0001, BF10 = 736571.5;SCZ mean vis–tac= 0.17, t(52)= 2.04,p= 0.093, BF10= 1.49). The difference between these two contrasts showed that HC had a larger difference between vis and tac conditions than SCZ (SCZ - HC m = 0.40, t(52) = 3.33, p = 0.002, BF10 = 21.00). Taken together, the analyses of ERPs to bisensory VT stimuli revealed aberrant long latency(> 230ms) IA effects in SCZ patients, suggesting an intersensory attention deficit in patients.

**Table 2:** Inter sensory At tention effects on ERPs to multisensory stimuli. Results for the main effects of t-test for each time-point and electrode of the full time-course of stimulus-evoked activity. Time windows were identified post-hoc by clustering analysis in order to structure the results. Amplitudes(μV)with in the cluster-algorithm defined TOI and ROI were then averaged. These averages were tested against Group and Attention in a follow-up linear mixed-model ANOVA to determine whether there were Group effects or Group*Attention interaction effects.

| Multisensory |  |  |  |  |
| --- | --- | --- | --- | --- |
| TOI | Cluster | $\beta$ | F(1,52) | p |
| 90-118ms | Group | 0.032 | 0.14 | 0.711 |
|  | attention | 0.18 | 48.43 | <.0001*** |
|  | Group*Attention | 0.03 | 0.40 | 0.530 |
| 158-216ms | Group | 0.16 | 1.10 | 0.299 |
|  | attention | 0.19 | 47.45 | <.0001*** |
|  | Group*Attention | -0.04 | 0.904 | 0.346 |
| 168-218ms | Group | -0.19 | 1.65 | 0.204 |
|  | attention | -0.23 | 42.94 | <.0001*** |
|  | Group*Attention | 0.04 | 0.48 | 0.493 |
| 230-320ms | Group | 0.27 | 1.72 | 0.196 |
|  | attention | 0.40 | 34.31 | <.0001*** |
|  | Group*Attention | -0.19 | 5.29 | 0.025* |
| 282-318ms | Group | -0.08 | 0.085 | 0.772 |
|  | attention | -0.34 | 38.60 | <.0001*** |
|  | Group*Attention | 0.21 | 11.60 | 0.002** |

*p* values: \*\*\* 0.001 \*\* 0.01 \* 0.05

**Figure 2:**
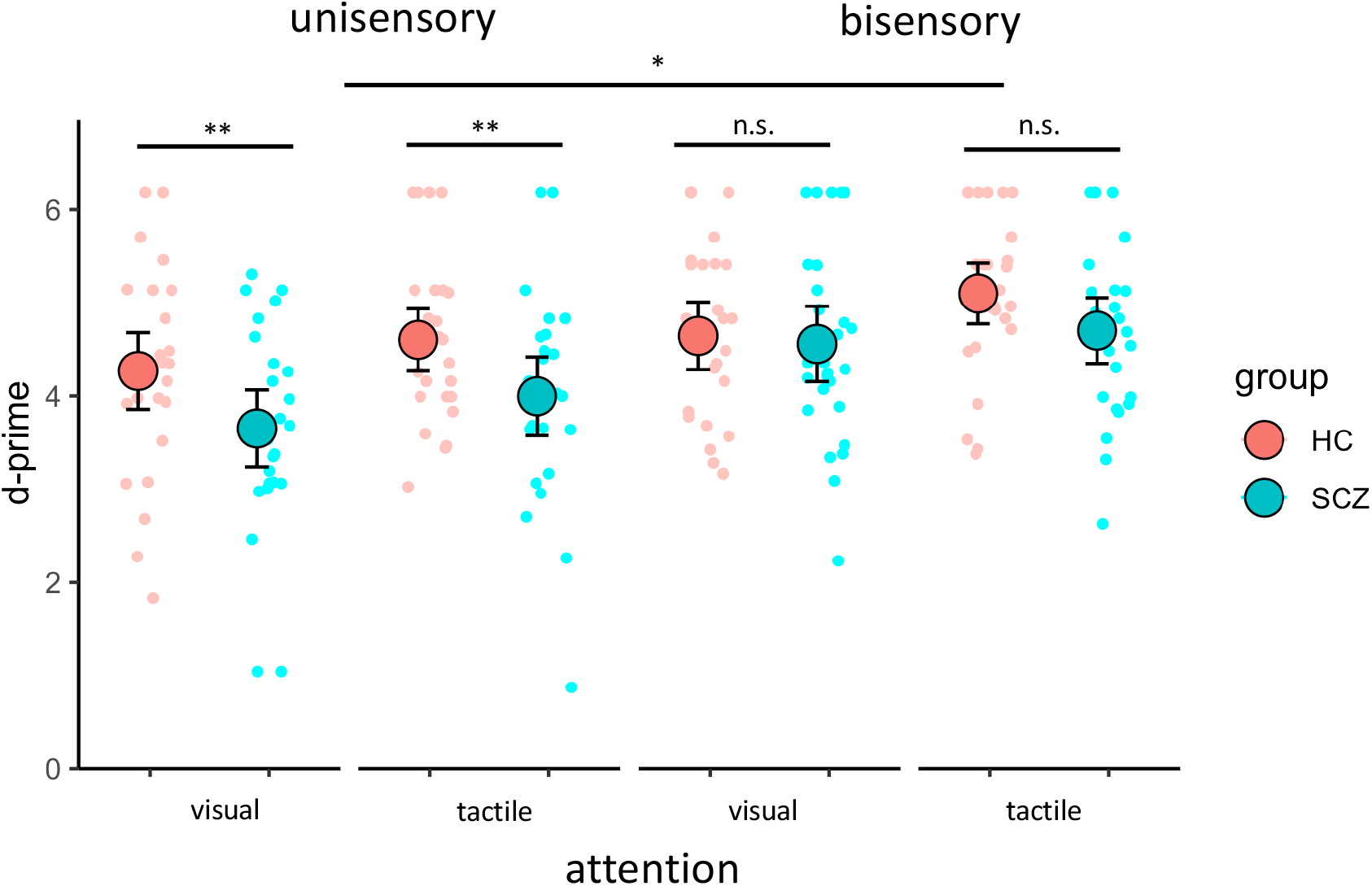
Behavioural data reveal intersensory attention deficits in schizophrenia for unisensory stimuli but not for bisensory stimuli. The figure shows d-prime values for unisensory and bisensory stimuli, visual and tactile attention conditions, and for the HC group (red points) and the SCZ group (turquoise points). Large points represent mean values, error bars represent ±1 SE, and the smaller points represent individual d-prime scores. Significant main effects and interactions are represented with asterisks (n.s., p > 0.05; * p < 0.05; ** p < 0.01). Across groups and attention conditions, the performance was better for bisensory compared to unisensory stimuli. Moreover, people with SCZ were worse that HC in the processing of unisensory visual and unisensory tactile targets. Notably, no such difference was found for bisensory targets, with Bayes Factor suggesting anecdotal evidence for the null hypothesis.

**Figure 3:**
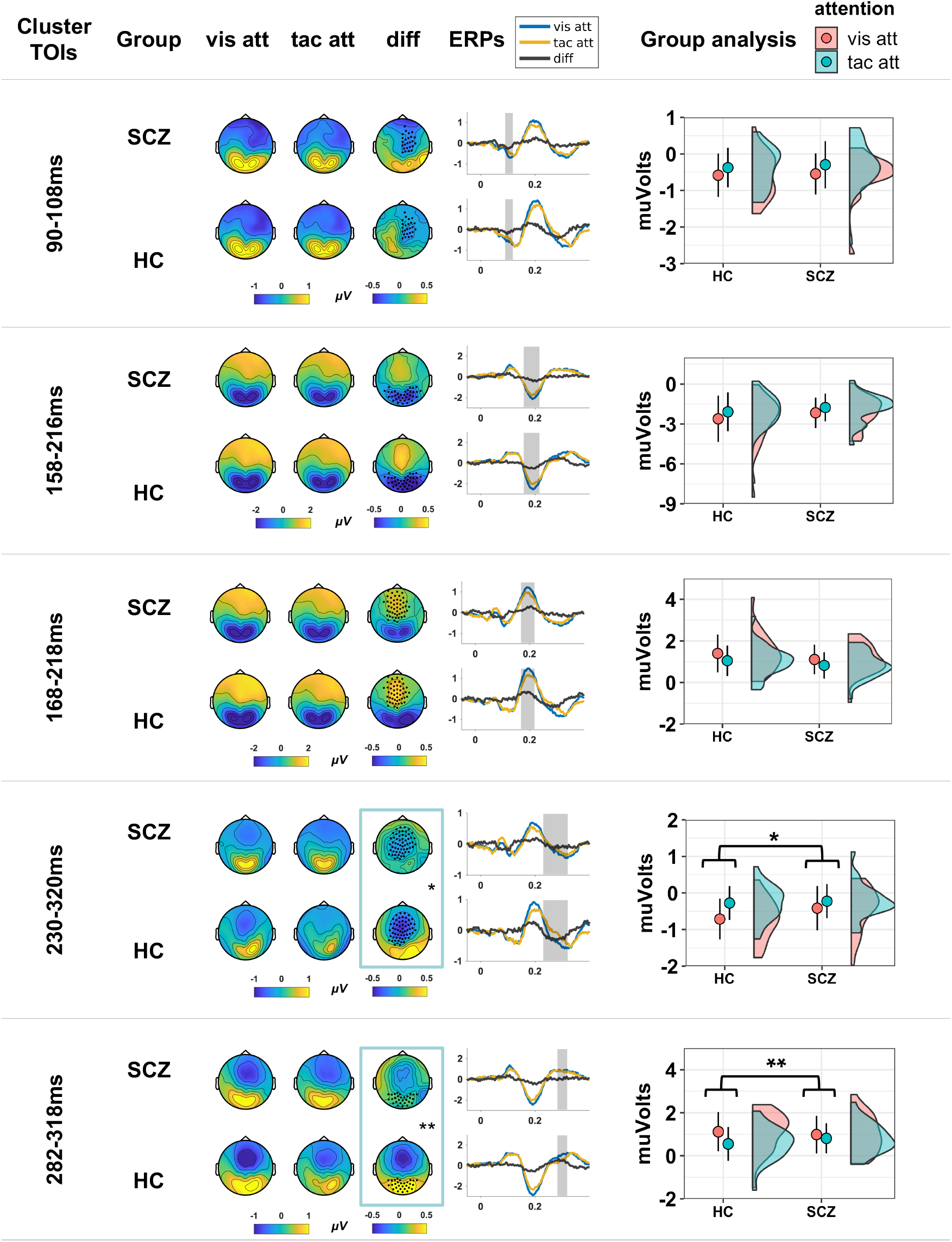
Intersensory attention effects in evoked brain potentials to bisensory VT stimuli are diminished in people with schizophrenia after 230ms. Topographic plots (left panel), traces (middle panel) and amplitude averages with confidence intervals (right panel) for five clusters of IA effects on bisensory VT stimuli. For the first three clusters (90ms to 218ms; upper three rows), IA effects were comparable between SCZ and HC groups. At longer latency (> 230ms; lower two rows), IA effects at mediofrontal and occipital scalp were significantly smaller in SCZ compared to HC. p-values: ** 0.01 * 0.05.

Finally, it the neural signatures of IA effects processing in the five clusters were correlated against the clinical PANSS and BACS scores as well as potential confound factors including nicotine consumption, and olanzapine equivalent dosage. These analyses revealed no significant relationships (all p-values > 0.05, uncorrected).

### Multisensory integration effects in evoked brain activity

The cluster analysis of MSI effects (bisensory VT vs. unisensory V+T)across groups revealed 4 significant clusters (Figure 4 and Table 3). There were two temporally overlapping clusters in occipital (240-280ms) and frontal (246-292ms) scalp regions, and two temporally overlapping clusters in frontal (316-364ms) and occipital (306-354ms) scalp regions. In the next step of the analysis, two-by-two factorial mixed model ANOVA (Group*Modality) were computed for each cluster. The ANOVAs revealed main effects of Group for the306-354ms and 316-364ms clusters, due to overall larger evoked brain responses in the HC compared to the SCZ group. More importantly, however, there were no group by Modality interactions in any cluster. For the frontal cluster at 316-364ms, both HC and SCZ showed significant differences between multisensory and additive conditions (HC multi - add mean difference = 0.914, t(52) = 5.42 <.0001, BF10 = 8815.24; SCZ multi - add mean difference = 0.675 t(52) = 4.00, p = 0.0004, BF10 = 120.85). Linear contrast of these differences were non-significant (SCZ – HC, m = 0.238, t(52) = 1.00, p = 0.323, BF10 = 0.41). For the posterior cluster at 306-354ms, both HC and SCZ showed significant differences between multisensory and additive conditions(HC multi - add mean difference = −1.15, t(52)= −6.06,p<.0001, BF10 = 71980.64; SCZ multi - add mean difference = −0.81, t(52) = −4.28, p = 0.0002, BF10 = 266.69). Linear contrast of these differences were non-significant (SCZ – HC, m = −0.34, t(52) = −1.257, p = 0.214, BF10 = 0.53). The Bayes Factor suggests anecdotal evidence for the null hypothesis in comparing HC and SCZ differences in MSI, indicating comparable multisensory integration effects in both groups.

**Table 3:** Multisensory integration effects on ERPs to bisensory vs. combined unisensory stimuli. Results for the main effects of t-test for each time-point and electrode of the fulltime-course of stimulus-evoked activity. Time windows were identified by clustering analysis in order to structure the results. Amplitudes(μV)with in the cluster-algorithm defined TOI and ROI were then averaged. These averages were tested against Group and Modality (bisensory vs. combined unisensory) in a follow-up linear mixed-model ANOVA to determine whether there were group effects or group*modality effects. MSI effects were found for both groups. The magnitude of the MSI effects did not significantly differ between groups, as shown by the absence of Group*Modality interactions.

| Multisensory vs. combined unisensory |  |  |  |  |
| --- | --- | --- | --- | --- |
| | Cluster | $\beta$ | F(1,52) | p |
| 246-292ms | Group | -0.06 | 0.27 | 0.60 |
|  | Modality | 0.21 | 50.69 | <.0001*** |
|  | Group*Modality | -0.01 | 0.03 | 0.86 |
| 184-236ms | Group | 0.22 | 2.51 | 0.12 |
|  | Modality | -0.18 | 47.40 | <.0001*** |
|  | Group* Modality | -0.03 | 0.32 | 0.57 |
| 316-364ms | Group | 0.21 | 4.54 | 0.04* |
|  | Modality | -0.35 | 44.34 | <.0001*** |
|  | Group* Modality | 0.08 | 1.00 | 0.32 |
| 306-354ms | Group | -0.24 | 6.27 | 0.02* |
|  | Modality | 0.38 | 53.49 | <.0001*** |
|  | Group* Modality | -0.10 | 1.58 | 0.21 |

*p*-values: \*\*\* 0.001 \*\* 0.01 \* 0.05

**Figure 4:**
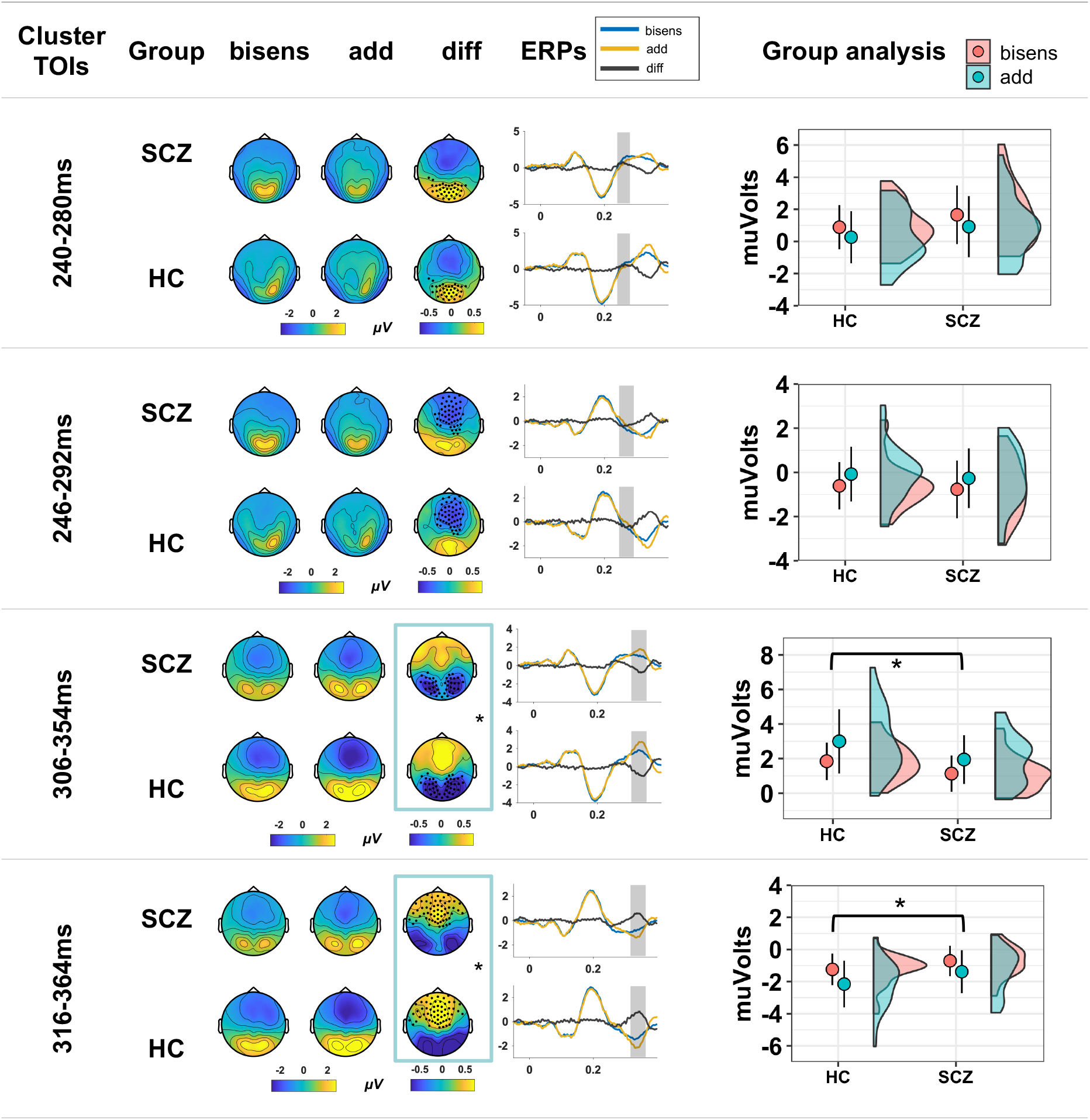
Multisensory integration effects do not differ between healthy controls and people with schizophrenia, suggesting intact MSI in patients. Topographic plots (left panel), traces (middle panel), and amplitude averages with confidence intervals (right panel) for the four significant clusters of MSI effects (i.e.bisensory VT vs. unisensory V+T) starting at 240ms. At the two later clusters (lower two rows), overall larger ERP amplitudes were found in the control group. p-values: * 0.05.

### Relationships between behavioural performance with intersensory attention and multisensory integration effects in schizophrenia

In this analysis, the relation between IA and MSI effects and d-prime behavioural performance in the SCZ group were tested. To this end, four Pearson correlations (i.e. clusters that showed main effects or interactions in relation to the factor Group) were calculated between bisensory d-prime values and two IA clusters(230-320ms,282-318ms)as well as two MSI clusters(306-354ms, 316-364ms). To account for multiple testing the alpha-level for the correlations was adjusted with Benjamini-Hochberg (α = 0.0375) (Benjamini & Hochberg, 1995). The analysis revealed that SCZ patients with larger IA effects in the 230-320ms cluster showed a better behavioural performance (230-320 ms; r = −0.47, p = 0.0138, Figure 5, left panel). Thus, patients with larger attention effects in the EEG data also performed better behaviourally. Moreover, the analysis revealed significant correlations between multisensory d-prime and the occipital cluster (306-354ms; r = −0.42, p = 0.030; 316-364ms Figure 5, middle panel) as well as the medial-frontal cluster (r = 0.40, p = 0.037, Figure 5, right panel). Hence, patients with larger MSI effects in the EEG data also performed better at the behavioural level. Taken together, the analyses show that IA and MSI mutually facilitate bisensory stimulus processing. It is possible that intact MSI compensates for attention deficits in patients.

**Figure 5:**
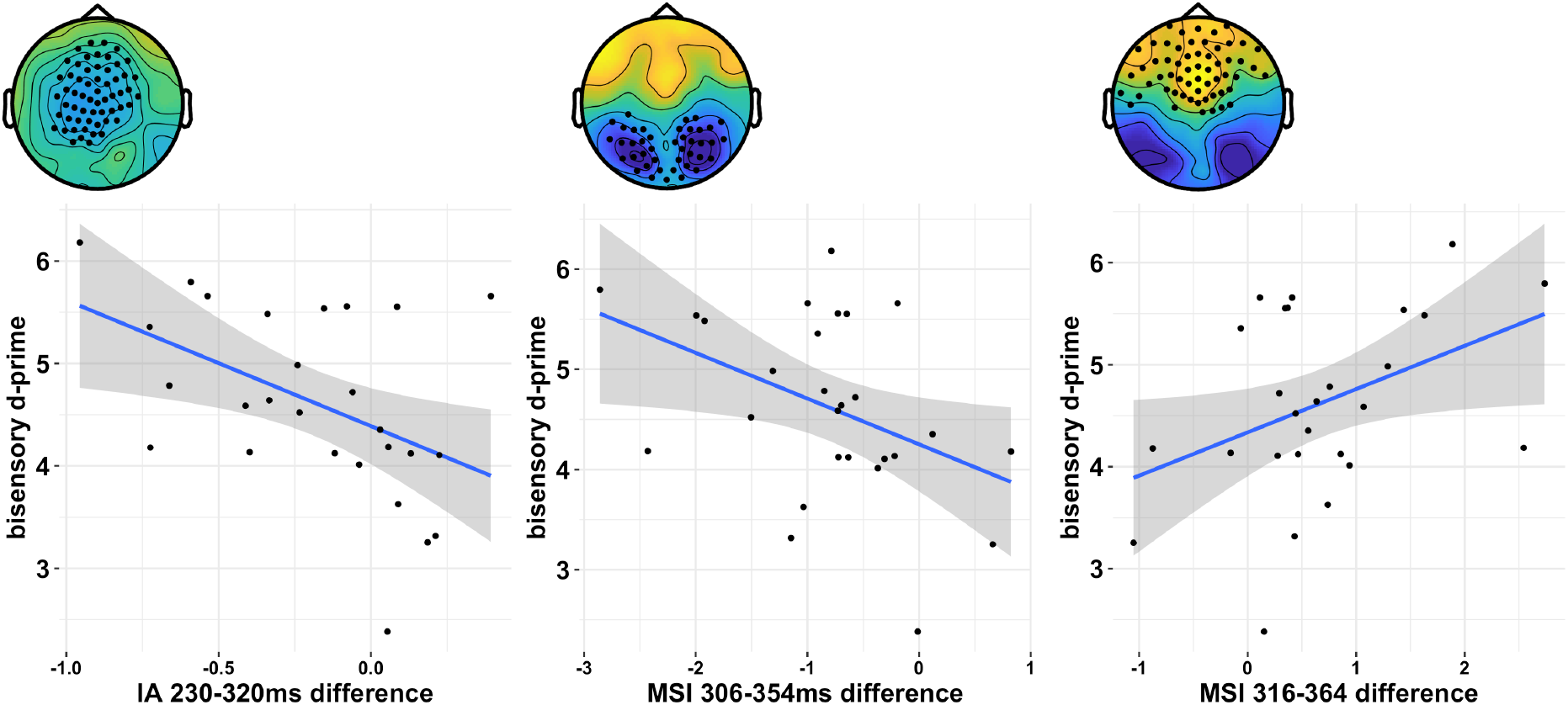
Scatterplots for SCZ group showing Pearson correlations multisensory d-prime against IA 230-320ms (left panel), MSI 306-354 (middle panel), MSI 316-364ms (right panel). In each case the graphs show that a better performance, i.e. higher d-prime for bisensory stimuli is associated with stronger IA and MSI effects in SCZ.

## Discussion

In this study we investigated the interplay between intersensory attention and multisensory integration in HC and in people with SCZ. People with SCZ showed deficits for unisensory target detection, however, they show ednormal behavioural performances for bisensory stimuli. The analyses of evoked electrical brain activity revealed diminished IA effects for people with SCZ over mediofrontal and occipital scalp regions at later processing stages (> 230ms). The analysis of MSI revealed multiple phases of integration, in which SCZ and HC showed comparable multisensory processing. Moreover, long latency IA and MSI effects were both positively related to the behavioural performance in patients. Collectively, our data suggest that intact MSI can compensate for aberrant top-down attention processing inSCZ.

When looking at the processing of bisensory visuo-tactile stimuli, as reflected in evoked brain activity, we found comparable attention effects for SCZ and HC up until around 200ms. We observed at tention effects over somato sensory regions at around 100ms and over visual regions at around 200ms, which corresponds to previous studies reporting IA effects for the N1 and P1 components (Keil et al., 2017; Lenartowicz et al., 2014; Talsma et al., 2009). Thus, our results suggest intact early IA effects in SCZ. Interestingly, later (> 230ms) ERP deflections showed more divergence of IA effects between groups. In the control group later IA effects included a medio-frontal negative deflection at around 275ms and a positive occipital deflection at around 300ms. Similar IA effects have been previously described in healthy individuals and have been interpreted as reflecting top-down processing (Eimer & Forster, 2003; Foxe et al., 2005; Gomez-Ramirez et al., 2016; Keil et al., 2017; Kida et al., 2004). In contrast to the HC group, there were significantly reduced long-latency attention effects in the SCZ group. Hence, our results suggest intact top-down attention effects on earlier sensory processing, but aberrant attention effects on later neural processing of bisensory stimuli in SCZ.

Using the additive approach to investigate multisensory processing for the same bisensory stimuli that were used for the analysis of IA effects, we found multiple phases of MSI. The first phase of MSI effects peaked at around 270 ms over occipital (around 260ms) and frontal (around 270ms) scalp regions, indicative for long-latency integrative processing. The lack of earlier MSI effects was somewhat surprising, since such effects have been reported previous studies in healthy individuals (Molholm et al., 2002; Talsma et al., 2007; Talsma&Woldorff, 2005). It is possible that our data-driven analysis approach, in which we corrected for multipletesting, haseliminated earlier MSI effects. Interestingly, the MSI effects in our study showed evidence of being comparable between groups, according to Bayes Factor calculation. This suggests intact multisensory processing in SCZ. General group deviations in more general stimulus processing manifested after 300ms, with two temporally overlapping later clusters in frontal (around 340ms) and occipital (around 330ms) scalp regions. In these later clusters, SCZ showed weaker evoked amplitude deflections for both multisensory and unisensory conditions. Diminished long latency ERP components in people with SCZ have been described in various previous attention paradigms (Michie et al., 1990; O’Donnell et al., 1999; Wood et al., 2007). However, although the amplitudes were generally smaller in SCZ, there were no significant differences in MSI effects between groups. Taken together, our data show multiple stages of MSI and suggest uncompromised multisensory processing inSCZ.

Of particular interest was our finding that the impairments in long latency attention processing at later latencies in occipital and mediofrontal regions were closely related to the behavioural performance in SCZ. This is interesting because patients showed normal behavioural performance for multisensory stimuli. Hence, it is possible that the additional information in multisensory stimuli compensates for the reduced long-latency attention deficit in SCZ. In other words, intact multisensory processes, which by themselves elevate and capture attention (Talsma et al., 2010), may boost the processing of bisensory stimuli in people with SCZ in a way that they are sufficiently attended to compensate for deficits in top-down attention processing. This claim is supported by our finding that MSI effects over overlapping late time windows with the attention effects were positive lycorrelated with the behavioural performance. Therefore, it is likely that the MSI effects, which occurred in parallel to the IA effects, compensate for long-latency attention deficits in SCZ.

Previous studies of MSI in SCZ are conflicted. Some studies have shown both impaired unisensory and multisensory processing in SCZ (Stevenson et al., 2017; Williams et al., 2010), where as others suggest that multisensory processing is relatively intact in patients (Stone et al., 2011; Wynn et al., 2014). Our findings align with the latter studies, in showing that SCZ have intact MSI. However, the additional manipulation of top-down attention in our study provides an important clue as to why previous research findings may have been conflicting. There is no obvious distinction between the tasks in these previous studies, which were basic, non-semantic multisensory tasks. Indeed, two employed similar target detection paradigms (Stone et al., 2011, and Williams et al. 2010), finding conflicting results. If attention was a factor in previous studies, it would be possible that the intrinsic attentional variability in different heterogeneous SCZ samples might have accounted for the different results. In support of this, Williams et al. (2010) did find a negative correlation between negative symptoms and MSI. Our own correlations between EEG IA and MSI effects against PANSS and BACS did not confirm any such relation. However, our experimental manipulation of attention provided a robust test of this influence in SCZ, which could be generalized to other MSI paradigms in future studies.

Our findings have some implications for theories of the aetiology of SCZ symptomology and its relation to sensory processing deficits. Researchers have suggested that the accumulation of processing errors cascades up into higher level distortions, resulting in positive and negative symptoms (Stevenson et al., 2017; Zhou et al., 2018). The observation of preserved MSI in SCZ suggests that perceptual processing deficits are not necessarily cumulative. Sensory information from a different modality can reduce error processing by providing statistically independent sampling from the environment (Gingras et al., 2009). It is possible that people with SCZ can use this to compensate for top-down attention deficits. Future studies could clarify this by systematically manipulating the reliability of individual cues in multisensory paradigms. Since multisensory processes can presumably compensate for top-down deficits in SCZ, we could further examine how patients respond to stimuli with variations in stimulus reliability. If input to one sensory modality was less reliable than input to another, could SCZ successfully assign corresponding weights to the incoming information according to the quality of the signal (Beauchamp et al., 2010)? Another theory of SCZ aetiology holds that distortions in coordination of perception could trigger limits in inner and outer boundaries (Postmes et al., 2014). Our findings suggest that attentional deficits in any failure of integration are an important part of such models. In sum, the aetiology of SCZ could be related to the interplay between MSI and top-down attention processes.

Our study had some limitations. We did not explicitly control for the length of time between measurement and onset of the first psychosis, so no relation between the progression of the disorder and IA or MSI could be tested. Schizophrenia is an extremely heterogeneous disorder and affected individuals have a wide range of symptoms and functional capacities. Alternative perspectives with people at early vs. later stages of SCZ, first-degree relatives and/or schizotypal personality disorder would help triangulate results. Nevertheless, our results provide a solid foundation for further investigation of the interaction between attention and multisensory integration at early stages of psychosis. Another limitation is the investigation of medicated patients. In the current study, we statistically controlled for medication and nicotine consumption, as well as excluding current those with current drugabuse, however these factors still contribute heterogeneity to the sample.

### Conclusion

Our study sought to examine the interplay between attention and multisensory processing in schizophrenia. At the behavioural level, we found attention deficits for unisensory stimuli, but not for multisensory stimuli. At the neural level, we observed aberrant attention processing of bisensory stimuli at longer latency over occipital and medio-frontal scalp regions, presumably reflecting aberrant top-down processing in SCZ. Our study suggests that multisensory processing, which appears to be intact in SCZ, can compensate for top-down processing deficits. It is possible that previous conflicting findings on multisensory processing in SCZ relate to differences in attentional demands across studies.

## Acknowledgements

This research was funded by a grant to DS from the German Research Foundation (SE1859/4-1).We thank Marianne Bröker, Teresa Ramme, Lisa Renziehausen, and Joseph Wooldridge for helping gather the data.

